# Testing the role of cognitive inhibition in physical endurance using high-definition transcranial direct current stimulation over the prefrontal cortex

**DOI:** 10.1101/566901

**Authors:** Gauthier Denis, Raphael Zory, Rémi Radel

## Abstract

The aim of this study was to clarify the role of the prefrontal cortex (PFC) in physical effort regulation. We hypothesized that the PFC would be progressively involved in physical endurance through the engagement of cognitive inhibition, which would be necessary to maintain effort by inhibiting fatigue-related cues. This hypothesis was examined using a double-blind, sham-controlled, within-subjects study (*N* = 20) using high-definition (HD) transcranial direct current stimulation (tDCS) over the dorsolateral prefrontal cortex (dlPFC). Participants had to maintain a knee extensor contraction at 30% of their maximal force while simultaneously performing an Eriksen flanker task to evaluate their inhibition performance during the task. Anodal stimulation of the dlPFC influenced response to the cognitive task during exercise, as seen by slower response times and better accuracy. However, it did not lead to any measureable improvement in cognitive inhibition and did not influence endurance time. There was no correlation between cognitive inhibition and the maintenance of physical effort. This result could be explained by some methodological limitations of our protocol, and we also provide alternative explanations for the contribution of the PFC in physical endurance.

## 1. Introduction

An increasing number of studies have tried to understand the brain’s involvement in the regulation of physical effort (Ekkekakis, 2009; McMorris, Barwood, & Corbett, 2018; Robertson & Marino, 2016; Tanaka & Watanabe, 2012). While fatigue of the musculoskeletal and cardiorespiratory systems certainly determines the cessation of effort, this relationship is indirect; effort termination is probably better reflected by the neural interpretation of signals of fatigue coming from the peripheral system. Accordingly, it has been shown that manipulating cognitive factors alone can result in different endurance times for a similar physical load (e.g., Ducrocq et al., 2017; Radel et al., 2017; see also McCormick et al., 2015 for a review). However, the neurocognitive mechanisms underlying the maintenance of physical effort remain largely unknown.

The primary motor cortex (PMC) has a well-known role in motor control and may therefore represent a candidate area for the maintenance of physical effort. A causal role of the PMC on endurance has been directly tested using transcranial direct current stimulation (tDCS), which can modulate the neuronal excitability of this region. The heterogeneity of the results means that there is no strong evidence that an increase in PMC excitability results in a higher capacity to endure physical effort (see Angius et al., 2017 for a review). Neuroimaging studies have highlighted the contribution of other cortical regions in the maintenance of physical effort. For example, a functional magnetic resonance imaging (fMRI) study indicated that, in addition to sensorimotor areas, frontal areas (the prefrontal cortex [PFC] and the anterior cingulate gyrus) were also engaged during a sustained contraction task (Liu et al., 2003). Near infrared spectroscopy (NIRS) studies have also consistently reported an increase of PFC activity during sustained exercise (see Rooks et al., 2010 for a meta-analysis). Because NIRS studies have also frequently observed a decline of PFC activity just before the cessation of exercise (Bhambhani et al., 2007; Rupp & Perrey, 2008; Tempest et al., 2014), it has been suggested that the PFC plays an important role in the maintenance of physical effort (Ekkekakis, 2009; Robertson & Marino, 2016). Traditionally, the PFC has been associated with cognitive rather than motor functions; thus, it has been suggested that PFC activity reflects the mobilization of inhibitory control mechanisms and, more precisely, cognitive inhibition (Ekkekakis, 2009; Perrey et al., 2016).

Cognitive inhibition is an executive function that relies heavily on prefrontal structures such as the inferior frontal cortex and dorsolateral PFC (dlPFC) (Aron, 2007). Cognitive inhibition is characterized by the blocking of automatically triggered processes when they prove to be unsuitable (Burle, Van den Wildenberg, & Ridderinkhof, 2005). More specifically, it allows individuals to suppress the impulses that compete with their voluntary goals. In the context of physical endurance, this might be seen as the conflict between an individual’s conscious goal to maintain effort/perform well and the peripheral signals of fatigue that lead to an impulse to stop. According to the hypothesis formulated by Ekkekakis (2009), as exercise intensity increases, cognitive mechanisms in the PFC may become active by exerting an inhibitory control over aversive stimuli to regulate the negative affective response. The maintenance of effort would therefore require the inhibition of painful physical sensations, a consequence of muscular activity during intense physical exercise (Cook, O’Connor, Eubanks, Smith, & Lee, 1997). This hypothesis is in line with several recent findings and theoretical propositions. For example, in a study that evaluated the cognitive determinants of pain sensitivity (Oosterman et al., 2010), only inhibition performance was associated with pain resistance. Specifically, better cognitive inhibition in a cognitive task was related to an increased immersion time and decreased pain sensitivity. Radel et al. (2016) proposed that the maintenance of ongoing physical exercise mainly relies on inhibition and a recent study (Cona et al., 2015) indicated that inhibition could distinguish the best ultra-marathon runners from the others, with a better cognitive inhibition performance for the best runners.

While previous work has therefore indicated there to be a relationship between cognitive inhibitory control and the maintenance of physical effort (Cona et al., 2015; Ekkekakis, 2009; Perrey et al., 2016), this has not yet been demonstrated experimentally. Thus, the objective of this study was to investigate the role of cognitive inhibition in physical endurance. To manipulate cognitive inhibition, tDCS was used to increase excitability of the PFC, as previous findings have successfully shown a tDCS-based modulation of inhibibition performance. Specifically, it was found that, compared to a sham condition, an anodal stimulation centered over the dlPFC led to an improvement in inhibition performance, as indexed by a decreased Stroop interference effect (Jeon & Han, 2012; Loftus et al., 2015) and a quicker post-conflict adjustment in the flanker task (Gbadeyan et al., 2016). In our study, high-definition tDCS (HD-tDCS) was used to attain a focal stimulation (Villamar et al., 2013) of the PFC and to avoid unwanted modulation of other regions. Anodal stimulation was centered around the right dlPFC, which is an area that has been linked to cognitive inhibition (Cipolotti et al., 2016). We hypothesized that active tDCS stimulation over the right dlPFC would result in better cognitive inhibition performance, and in turn, lead to better endurance performance compared to a sham tDCS condition. The Eriksen flanker task was implemented during the physical task to measure inhibition performance. We therefore expected that cognitive inhibition would play a mediating role that would explain how modulation of PFC activity can influence endurance time.

## 2. Material and methods

### 2.1. Participants

In line with previous studies that have reported a moderate-to-large size effect of anodal tDCS on inhibition (Gbadeyan et al., 2016; Jeon & Han, 2012; Loftus et al., 2015), we estimated that at least 19 participants would be required to find a significant effect (alpha=.05) of this magnitude (*d*=.60) with an 80% chance level. Accordingly, 20 right-handed healthy adults participated in the study (7 female, 13 male, 20.55±1.73 years old). Participants were students of a French university who participated in exchange for course credits. Participants signed a consent form to participate in the study but remained unaware of the specific research hypotheses. To respect safety recommendations for the use of tDCS (Woods et al., 2016), exclusion criteria were the presence of metal in the head, a pacemaker, or brain injury.

### 2.2. Procedure

Participants were invited to the laboratory on two separate occasions that were held at the same time of day. To eliminate the effects of residual fatigue, the two sessions were separated by a minimum of four days. Participants were instructed to abstain from any vigorous exercise for 24 hours before the session, and to sleep at least 7 hours the night before. Apart from the type of stimulation (anodal tDCS over the right dlPFC or sham stimulation), all sessions were identical (Figure 1). For each session, participants were first equipped with electromyography (EMG) electrodes on the quadriceps and tDCS electrodes on the scalp. The participants were then seated in a dynamometric chair with the right leg (right ankle) fixed on a padded support that allowed the knee to form an angle of 90 degrees. Learning of the cognitive task was then implemented, whereby participants had to complete four 2-min blocks of the Eriksen flanker task. Additional blocks were carried out if the learning criteria were not achieved, including a between-block performance variability under 5%, an average reaction time (RT) that was less than 550 ms, and a response accuracy above 85%. In the second experimental session, participants completed the same number of task blocks to control for the potential effects of cognitive fatigue.

**Figure 1.**
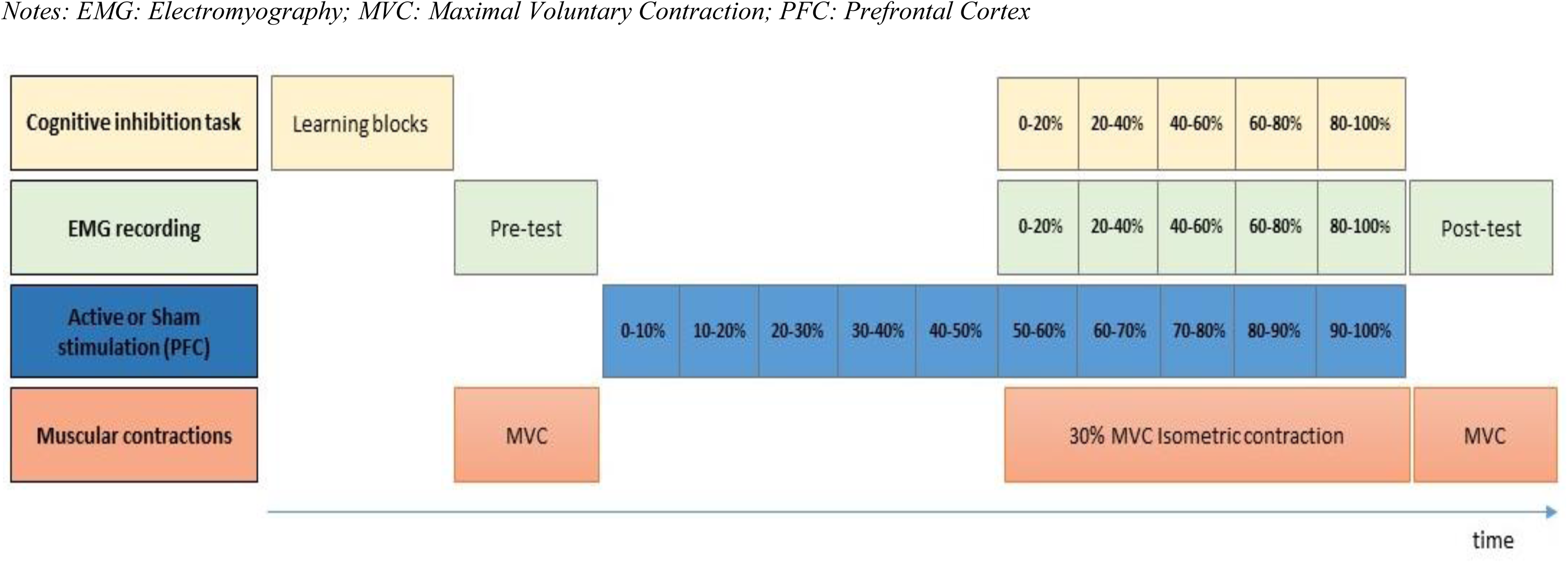
Illustration of the protocol of an experimental session

The motor task required participants to exert an upward-directed force with their right leg using the quadriceps (Angius et al., 2016). To determine the maximal voluntary contraction (MVC) force, participants were first instructed to exert a maximal force for 3 sec. Each MVC measure included two trials separated by a 60 s rest period. For each session, the largest peak value was retained as the MVC force and used to calculate the target force in the endurance task. Immediately after the MVC measurement, the tDCS stimulation conditions began in a double-blinded fashion. The first 10 min of stimulation was applied to increase the chances of modulating PFC activity during the upcoming task. Stimulation was maintained during the endurance task. The aim of this stimulation design was to facilitate cognitive inhibition, which would in turn lead to better endurance performance in the active stimulation condition (Hogeveen et al., 2016). By pulling a wire strapped to their ankle, participants had to exert the force required to hold a weight above the ground through a pulley system. After accounting for all friction forces, the necessary force to lift the weight corresponded to 30% of their MVC. The task was terminated when the participant was no longer able to maintain the weight above the ground. When the weight was close to the ground, participants were warned by a contact between their calves and an elastic band. In parallel with this sustained contraction task, participants had to perform the Eriksen flanker task. The cognitive task continued for as long as the participant was able to maintain the contraction. Immediately after stopping the task, the MVC force was assessed again in the same way as in the first measurement. Participants were then asked to evaluate their rating of perceived exertion (RPE) using a visual analogue scale (VAS). The use of a VAS avoids memorization of responses from one session to the next (Grant et al., 1999).

#### 2.2.1. Force recording

The signal from the force transducer was sampled at 100 Hz using an MP100 acquisition unit and the AcqKnowledge software (BIOPAC, Goleta, CA, USA). The dynamometric chair measured knee extension force using a torque transducer (SM 2000N, Interface, Scottsdale, USA). The pre-post task MVC difference was calculated to measure the extent of fatigue.

#### 2.2.2. EMG recording

EMG activity was recorded using 8-mm Ag/AgCl surface electrodes placed over the *vastus lateralis* (VL). EMG was recorded continuously during the whole experimental session using a BIOPAC MP100 system and AcqKnowledge software. The signal was filtered (3.5 Hz to 350 Hz), pre-amplified (×1000), and sampled at 2 kHz. For the MVC, the root mean square (RMS) of EMG activity was calculated over a 500-ms period around the highest MVC value (i.e., 250 ms before and 250 ms after the peak MVC). During the endurance task, RMS was calculated over five-time periods (corresponding to 0-20, 20-40, 40-60, 60-80, and 80-100% of the total duration).

#### 2.2.3. tDCS manipulation

Direct current stimulation was delivered using a StarStim wireless neurostimulator (Neuroelectrics, Barcelona, Spain). A 4×1 HD-tDCS montage was used to provide focal stimulation (Villamar et al., 2013). The central anode electrode was located as close as possible to the coordinates corresponding to the center of mass of the right dlPFC (MNI: x = 40, y = 14, z = 28) during the cognitive inhibition tasks (Nee et al., 2007). The real localization was obtained using 3D digitization (Patriot, Polhemus, Colchester, VT, USA). The four cathode electrodes were located at a distance of 3.5 cm around the anode electrode to ensure that the current would reach the cortical layer while maintaining a focal stimulation. Figure 2 illustrates the simulation of the electric field distribution for this montage using the StimWeaver option of NIC software (version 2.0, Neuroelectrics, Barcelona, Spain). The set of electrodes was held in place using a thermoplastic curved plate connected to elastic bands that were attached around the head. Following recommendations to optimize comfort and safety, Ag/AgCl sintered ring electrodes were used (Minhas et al., 2010) with a conductive gel. Impedance of each electrode was kept under 5 kOhm. If the impedance exceeded this limit during testing, the experimental session was cancelled and participants were asked to reschedule the session for another time. Current intensity was set at 2 mA in all HD-tDCS sessions. Depending on the experimental sessions, participants received either an anodal stimulation over the right dlPFC or a sham session (over the right dlPFC). The double-blind mode of the NIC software was used to ensure that neither the experimenter nor the participants knew if the session included real or sham stimulation. Each session file was prepared in advance by an independent person and the name of the file launched by the experimenter did not contain any identifying information about the condition. The order of the sessions was randomized and counterbalanced across participants. In each session, the stimulation began 10 min before the task and was maintained during the entire task (Figure 1). For the sham condition, the electrical current was only applied during the first and last 30 s of stimulation to induce the same cutaneous sensation as real stimulation.

**Figure 2.**
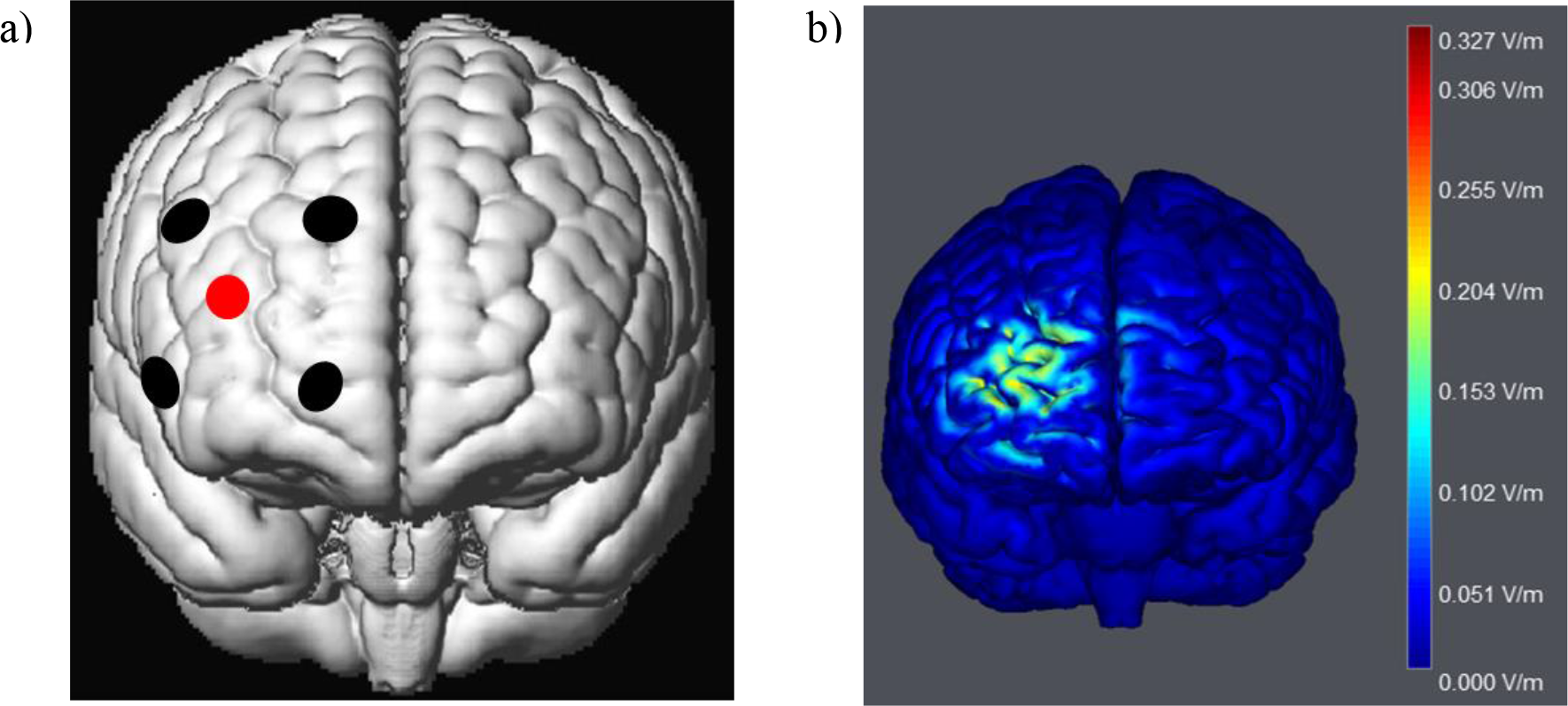
Cortical projection of the tDCS electrodes (one anode in red and four surrounding electrodes for the current return in black). a) representation of the montage for active or sham stimulation of the prefrontal cortex; b) representation of the simulation of the electric field distribution on the gray matter surface (in V/m) elicited by the anodal stimulation of the prefrontal cortex.

#### 2.2.4. Cognitive task

Cognitive inhibition was measured using a version of the Eriksen flanker task (Eriksen & Eriksen, 1974). Each trial began with the presentation of a fixation cross at the center of the screen. After 200 ms, the stimulus was presented and participants had to respond according to the direction indicated by the central arrow. Delivery of the response turned off the stimulus. When participants failed to respond within 1500 ms, the trial was terminated and the next trial began immediately. The task comprised two types of trials occurring with the same probability, as follows: congruent (CO) trials (all arrows were uni-directional) and incongruent (IN) trials (50% of central arrows were contra-directional). In this version, each group of arrows were randomly displayed at the top or at the bottom of the screen (Schmit et al., 2015) to prevent participants to focus their gaze on one point of the screen in order to avoid the flankers. Performance was calculated for five time periods (corresponding to 0-20, 20-40, 40-60, 60-80, and 80-100% of the total duration). RT and accuracy (% correct) of each type of stimuli were recorded. In addition, a number of other inhibition indexes were considered. The flanker effect, which represents the time required to resolve the interference, was obtained by subtracting RT in IN trials from RT in CO trials. We also looked at this flanker effect in the 20% of the longest trials, as these trials reflect more inhibition as this cognitive function is a slow and controlled process that takes time to build up (Ridderinkhof et al., 2004). Because an effect of tDCS on conflict adaptation has been observed in previous work (Gbadeyan et al., 2016), we also analyzed modulation of the flanker effect as a function of the type of preceding trial.

#### 2.2.5. Data analysis

All results were screened for extreme values using the outlier labelling rule (Hoaglin & Iglewicz, 1987), and extreme values were removed when detected. The distribution of all variables was visually inspected and checked using the Shapiro-Wilk and Kolmogorov-Smirnov tests. Linear mixed models (LMM) were used to analyze normally distributed data, and generalized linear mixed models (GLMM) were used non-normal data. If no known distributions could be fitted to the data for the GLMM, LMM were used after normalizing the data using the Box-Cox method (Box & Cox, 1964; Osborne, 2010). LMM or GLMM were used for statistical testing because they can attain a higher level of statistical power than traditional repeated measures analysis of variance (Ma et al., 2013) and are also recommended to prevent type 1 errors (Boisgontier & Cheval, 2016). A random intercepts effect structured by participants was included to control for the non-independence of the data and inter-subject variability. In line with recent recommendations (Barr et al., 2013), models with and without inclusion of a random slopes effect of the tDCS condition were compared using the Akaike information criterion, and the model with the smallest Akaike information criterion was retained.

The LMM for the analysis of endurance time included the order of the experimental sessions (to control for order effects) and tDCS condition (active vs. sham stimulation) as fixed factors. In addition to these factors, a fixed factor representing the measurement time was included in the LMM for the analysis of EMG and cognitive performance results. A factor representing the type of the preceding trial (CO vs. IN) was included in the model on conflict adaptation. For each dependent variable, Cohen’s *d*s were calculated to represent the effect size of the condition effect using the difference of the exact means and standard deviations between the HD-tDCS and sham sessions.

To identify a possible mediation effect of tDCS on endurance time via cognitive inhibition, we first explored the correlation between indexes of inhibition (the Flanker effect) and endurance time. The mediation hypothesis was then specifically tested using the PROCESS toolbox (Hayes, 2012). Condition (right dlPFC HD-tDCS or sham stimulation) was used as the independent variable, the Flanker effect as the mediator, and session order as a covariate for the prediction of the dependent variable (endurance time).

## 3. Results

### 3.1. Endurance time

The LMM for endurance time (without random slopes of the condition) revealed no significant main effect of stimulation condition [*F*(1,18) = 1.175, *p* = .293], whereby there was no significant difference between the sham (226.967 ± 101.275 s) and active (251.826 ± 98.330 s) stimulation conditions (Figure 3), and a small effect size (*d* = 0.25).

**Figure 3.**
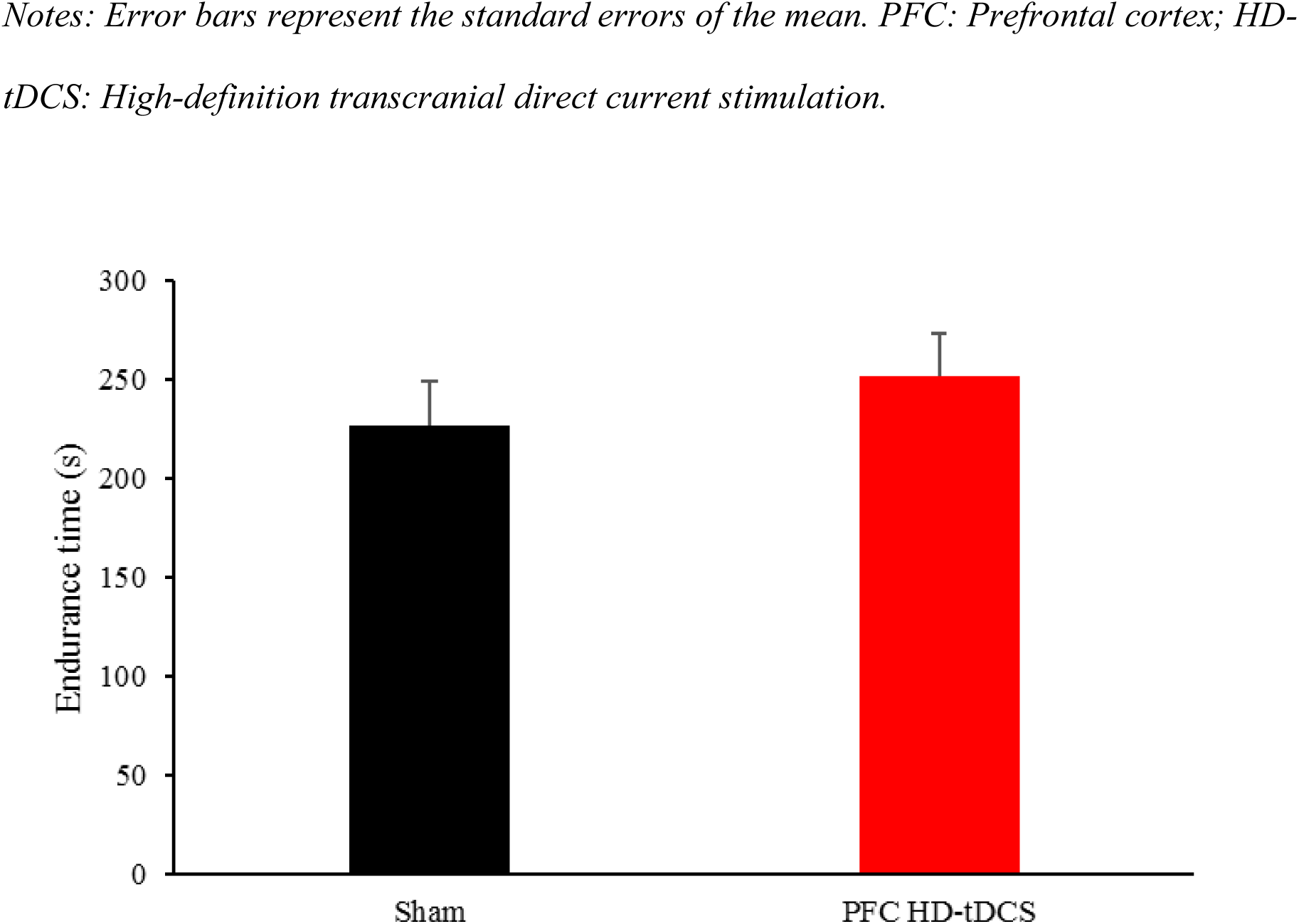
Endurance time as a function of the tDCS condition (sham or HD-tDCS, centered over the PFC)

### 3.2. Rating of perceived exertion

The LMM for RPE (without random slopes of the condition) revealed no significant main effect of stimulation [*F*(1,18) = 0.006, *p* = 0.941, *d* < 0.1].

### 3.3. Force recording

The LMM for MVC force (without random slopes of the condition) revealed a significant main effect of the time of measurement [*F*(1,56) = 46,940, *p* < 0.001], whereby there was a decrease in MVC force from the pre-test (49.962 ± 9.546 K/g) to the post-test (45.923± 10.05 K/g). There was no significant main effect of stimulation condition on MVC force [F(1,56) = 0.168, *p* = .684, *d*<0.1] and no interaction [F(1,56) = 0.072, *p* = .789].

### 3.4. Electromyography

#### 3.4.1. During MVC

The EMG-RMS data of the VL were normalized using a Box-Cox transformation. The LMM (without random slopes of the condition) revealed no significant effect of the time of measurement from the pre-test to the post-test [*F*(1,56) = 2.963, *p* = .091] or of the stimulation condition [*F*(1,56) = 0.695, *p* = 0.408, *d* < 0.1], and no interaction [*F*(1,56) = 0.122, *p* =.728].

#### 3.4.2. During sustained contraction task

The EMG-RMS data of the VL were normalized using a Box-Cox transformation. The LMM for EMG-RMS (with random slopes of the condition) revealed a significant effect of the time of measurement [*F*(1,157.025) = 94.393, *p* < 0.001], whereby VL activity increased over time in both conditions. However, there was no significant main effect of stimulation condition [*F*(1,24.825) = 3.547, *p* = .071, *d* < 0.1] and no interaction [*F*(1,157.025) = 2.073, *p* =.152].

### 3.5. Cognitive task

#### 3.5.1. Response time

Over the total duration of the task (239.396 ± 99.326 s), RTs were calculated for correct responses only (6894 trials analyzed), excluding the first trial (start of the task), the last trial (end of the task), and quick responses (RTs less than 100 ms), which were considered as anticipations. The GLMM on RT data was modelled with a gamma function (Lo & Andrews, 2015). The GLMM (without random slopes of the condition) revealed a main effect of the type of trial on RTs, whereby subjects had slower RTs in IN trials than in CON trials (482.10 ms vs. 553.86 ms). However, this factor also interacted with the time of measurement (treated as a continuous variable) [*F*(1, 6885) = 10.913, *p* = .001]. While RT decreased over time for compatible trials [*F*(4, 3950) = 1.882, *p* = .111], it increased for incompatible trials [*F*(4, 2931) = 3.948, *p* = .003]. A main effect of stimulation condition on RTs was found [*F*(1, 6885) = 5.991, *p* = .014, *d* < 0.1], whereby participants responded faster in the sham condition than in the dlPFC stimulation condition (508.83 ms vs. 511.27 ms). Interestingly, the stimulation condition also interacted withthe time of measurement [*F*(1, 6885) = 5.509, *p* = .019; Figure 4]; RT increased over time in the sham condition [*F*(4, 3322) = 2.958, *p* = .019], but remained stable in the dlPFC stimulation condition [*F*(4, 3558) = 1.001, *p* = .406].

**Figure 4.**
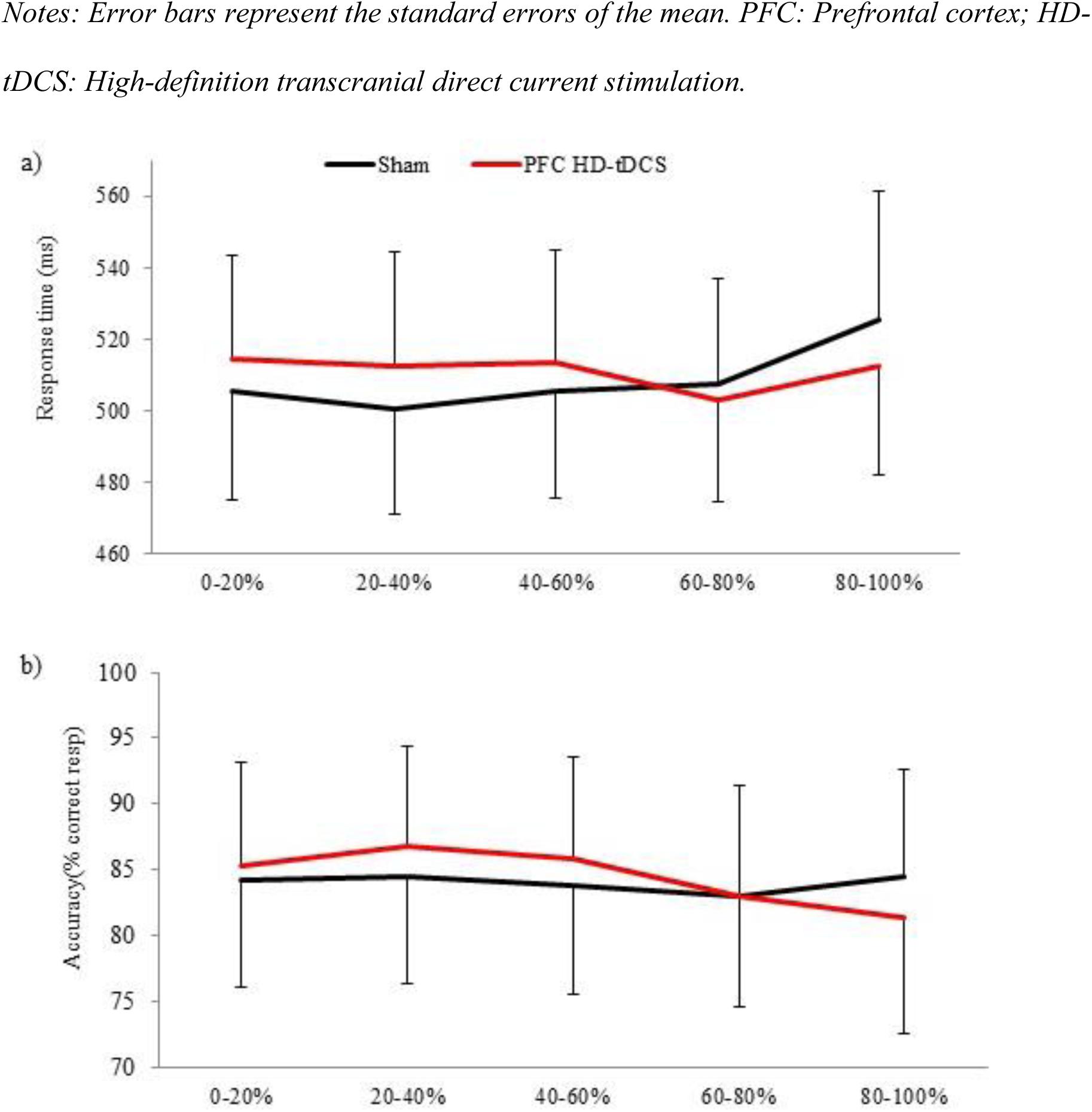
Response time (a) and accuracy (b) as a function of the tDCS condition (sham or HD-tDCS, centered over the PFC)

#### 3.5.2. Accuracy

The analysis of accuracy included all trials except the first trial, the last trial, and trials with a RT less than 100 ms (8184 trials analyzed). The GLMM on accuracy was modelled with a logistic function to account for binary data. The GLMM (without random slopes of the condition) showed a main effect of the type of trial [*F*(1, 8175) = 71.490, *p* = .000], whereby there was a lower accuracy for incompatible trials than for compatible trials (71.9% vs. 96.4% accuracy, respectively). There was an interaction effect between the type of trial and the time of measurement [*F*(1,8175) = 10.968, *p* = .001]. While accuracy increased over time in compatible trials [*F*(4,4091) = 0.539, *p* = .707], it deteriorated over time for incompatible trials [*F*(4,4071) = 3.554, *p* = .007]. There was a significant main effect of stimulation condition on accuracy [*F*(1,8175) = 4.039, *p* =.044, *d* = 0.28], whereby participants responded more accurately in the dlPFC stimulation condition than in the sham condition (84.4% vs. 84.0%). There was also an interaction between the stimulation condition and the time of measurement [*F*(1,8175) = 4.039, *p* < .05; Figure 3]. While accuracy remained stable during the task in the sham condition [*F*(4,3955) = 0.307, *p* = .873], it deteriorated over time in the dlPFC stimulation condition [*F*(4,4215) = 4.479, *p* =.001].

#### 3.5.3. Measures of inhibition

For the Flanker effect, the LMM for delta (without random slopes of the condition) revealed no significant main effect of stimulation condition (*d* = 0.12) and no interaction between stimulation condition and the time of measurement (*p*s >.05). For the Flanker effect at long RTs, the LMM (without random slopes of the condition) revealed no significant main effect of condition (*d* < 0.1) and no interaction between the condition and the time of measurement (*p*s > .05). Concerning conflict adaptation, the GLMM (without random slopes of the condition) revealed a significant effect of the preceding trial type on RT [*F*(1,6812) = 8.547, *p* = .003], whereby RTs were longer after an incompatible trial than after a compatible trial (510.55 vs. 519.96 ms). There was a significant interaction between the type of trial and the preceding trial accuracy [*F*(1,6812) =18.265, *p* = .000]; RTs increased for incompatible trials when the preceding trials were correct [*F*(4,5742) = 383.701, *p* = .000] or incorrect [*F*(1,1066) = 56.504, *p* = .000].

#### 3.5.4. Relationship between endurance time and cognitive inhibition

The results indicated that endurance time was not associated with the Flanker effect (*r* = .007, *p* = .967) or with the Flanker effect at long RTs (*r* = -.034, *p* = .833; Figure 5). The mediation model was not supported because the preliminary assumptions were not met; the independent variable had no effect on the mediator and on the dependent variables, and the mediator also had no effect on the dependent variable.

**Figure 5.**
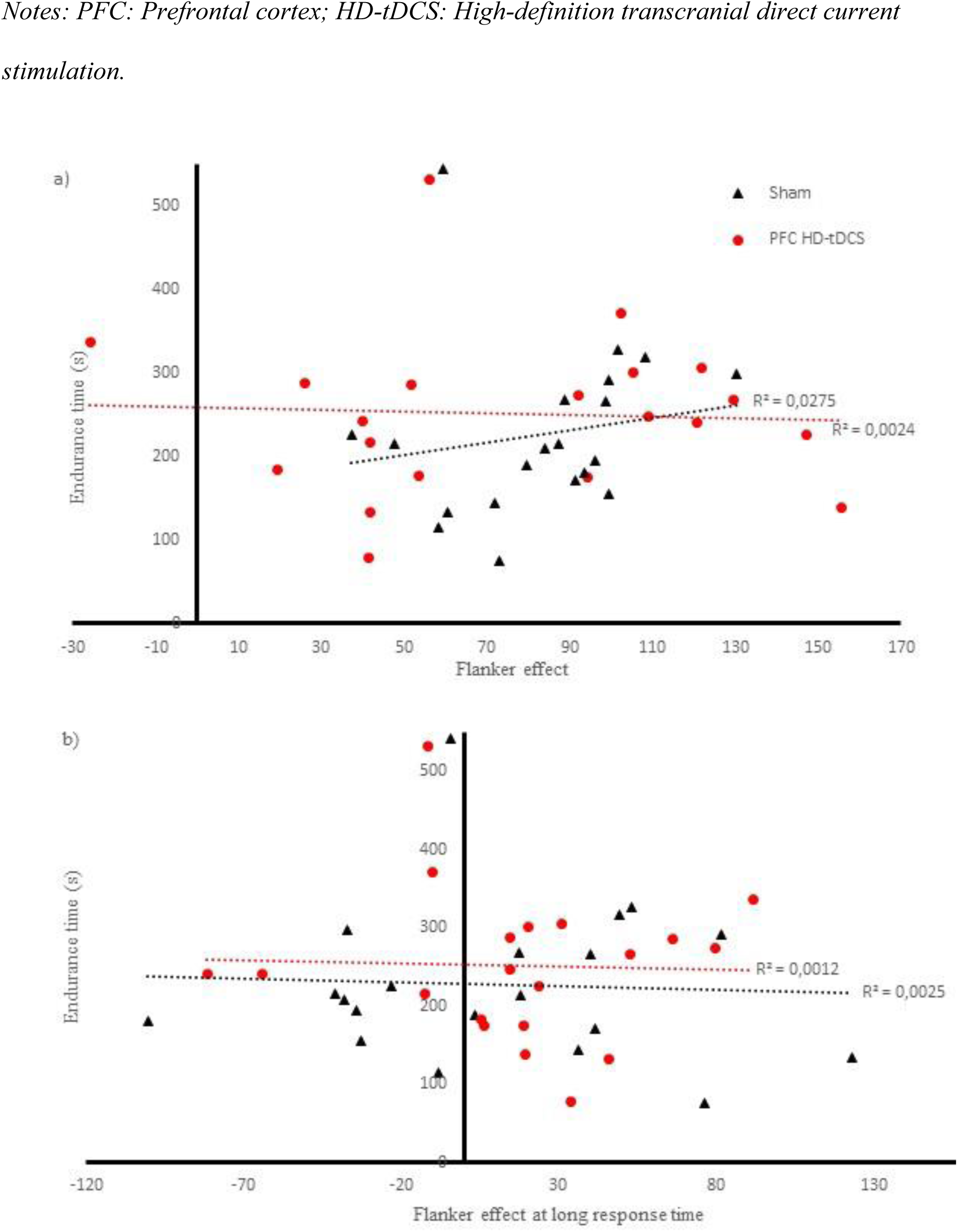
Correlation between cognitive inhibition and physical endurance. a) the correlation between endurance time and the flanker effect; b) the correlation between endurance time and the flanker effect for longer response times.

## 4. Discussion

This study examined the role of cognitive inhibition on the maintenance of physical effort. Our hypothesis was that facilitation of PFC activity would improve the capacity for cognitive inhibition and, in turn, delay the cessation of effort. Our results indicated that if the physical task successfully generated physical fatigue (increased the EMG signal throughout the task and decreased maximal force after the task), the electrical stimulation of the right dlPFC did not lead to any increase in endurance time. Furthermore, stimulation of the right dlPFC did not affect cognitive inhibition performance. Finally, we found no correlation between the maintenance of physical effort and cognitive inhibition. These observations are discussed separately below.

### 4.1. The impact of tDCS on endurance time

Contrary to our hypothesis, the use of anodal stimulation over the PFC did not lead to longer endurance times. However, it should be noted that similar null results have been reported by previous studies that attempted to influence physical endurance by modulating cortical excitability using tDCS (Kan et al., 2013; Muthalib et al., 2013; Radel et al., 2017; Williams et al., 2013). Despite these results, recent reviews have indicated that the net effect resulting from all these previous studies was nonetheless positive, and of a moderate amplitude (Angius et al., 2017; Radel et al., 2017). That said, the reliability of many previous studies is questionable due to a lack of double-blinding of experimental conditions and small sample sizes. We used double-blinding and an HD stimulation technique to provide better methodological control than conventional stimulation (Caparelli-Daquer et al., 2012). Most previous studies that have examined the effect of tDCS on physical endurance were also limited by the use of non-focal stimulation. For example, a recent study observed a significant improvement in endurance time with tDCS over the motor cortex, but concluded that this improvement was not due to changes in motor cortex excitability (Abdelmoula et al., 2016). The authors hypothesized that the use of non-focal tDCS may have influenced the activity of other cortical areas, which could have been responsible for the longer endurance time. These results lead us to believe that other regions benefit from the tDCS more than the motor cortex thus highlighting the importance to use a focal stimulation to ensure a precise determination of the neural structure responsible for the effect.

Following the proposition by Robertson and Marino (2016) about the implication of the PFC in the maintenance of physical effort, our focal stimulation was centered over the right dlPFC. Only one previous study has tested the effect of an anodal HD-tDCS stimulation centered on the right PFC, and similar non-significant results were reported (Radel et al., 2017). If an effect exists, it is possible that it would be of a small amplitude and could therefore not be observed in these studies as doing so would require a very high level of power. It should also be noted that there is now a growing consensus for the presence of variability in the individual responses to non-invasive brain stimulation (Guerra et al., 2017), and to HD-tDCS specifically (Wiethoff et al., 2014). However, even if we accept the interpretation that an increase in excitability of the right dlPFC does not modulate physical endurance, this does not entirely rule out the potential contribution of the PFC during endurance tasks. Other regions of the PFC that were not targeted by our HD-tDCS manipulation (e.g., the medial frontal cortex, inferior frontal cortex, and the left lateral PFC) might still contribute to the maintenance of the physical effort. For example, because endurance performance also reportedly depends on decision making (McMorris et al., 2018; Robertson & Marino, 2016), the medial PFC might play an important role through its involvement in the evaluation of costs (pain, fatigue, effort) and benefits (objectives, goals, rewards) of a given task (Domenech & Koechlin 2015; Kringelbach, 2005) With this in mind, it would be interesting to determine if modulation of orbitofrontal cortex activity would modify endurance performance through the alteration of decisional processes.

### 4.2. Impact of tDCS on cognitive inhibition

We found an effect of the HD-tDCS manipulation on general cognitive behavior during the Eriksen flanker task. The focal tDCS manipulation over the right PFC therefore successfully modulated cognitive activity. Specifically, participants respond slower and more accurate in anodal stimulation condition. The speed/accuracy trade-off also evolved differently throughout the task depending on the stimulation conditions as indexed by slower RTs and a stable accuracy over time in the sham condition, while the anodal stimulation led to a decline in accuracy but no RT slowdown over time (Figure 4). This result corresponds to results from recent systematic reviews and meta-analysis, which also showed that anodal tDCS over the dlPFC influences RTs and accuracy on cognitive tasks (Dedoncker et al., 2016; Hill, Fitzgerald, & Hoy, 2016).

However, besides these strategic effects on the task response, we found no clear changes on cognitive inhibition. We derived different metrics to reflect cognitive inhibition (i.e., the Flanker effect to represent the time taken to solve the interference; the Flanker effect at longer RTs, for which inhibition is usually more visible; and conflict adaptation to represent adjustment after incompatible trials) but failed to observe any effects of stimulation condition on any of these measures. This finding contrasts with previous reports that tDCS stimulation of the lateral PFC improved conflict adaptation during a flanker task (Gbadeyan et al., 2016) and reduced the time needed to solve interference (Jeon & Han, 2012; Loftus et al., 2015). It is difficult to explain the origins of these different results. The inter- and intra-individual variability in response to tDCS stimulation could be a candidate, as high variability has been observed not only in the motor domain but also in cognitive responses (Summers et al., 2016).

### 4.3. Relationship between cognitive inhibition and physical endurance

An important objective of the present study was to test the role of cognitive inhibition on physical endurance. For this reason, we also looked at the correlation between the different inhibition indexes and endurance time but found no significant correlations. The effect size suggests that this absence of correlation is not likely to be due to a lack of statistical power. We are only aware of one study that tested the relationship between cognitive function and endurance performance. Unlike the present study, a significant correlation between inhibition capacity and endurance performance was found (Cona et al., 2015). Similarly, a previous study highlighted that cognitive inhibition was associated with pain resistance (Oosterman et al., 2010). Since physical effort may cause painful sensations, these results suggest a role for cognitive inhibition performance in endurance performance. Besides cognitive inhibition, other studies have shown that attention might play an important role in endurance performance (Brick et al., 2016a, 2016b; Masters & Ogles, 1998). Because inhibitory processes are required to adopt an effective attentional strategy (Hasher et al., 2007), these results provide indirect support for the role of inhibition in endurance performance.

There are several potential reasons for the differences between previous findings and our own, which failed to find a significant correlation between cognitive inhibition and endurance. First, inhibition is not always treated as a unitary concept, and some models distinguish cognitive inhibition from response inhibition and behavioral inhibition (Howard et al., 2014). According to this view, cognitive inhibition concerns the suppression of a mental process, whereas response inhibition rather reflects the suppression of an automatically activated response or a behavioral impulse. The positive correlation between inhibition and endurance found by Cona et al. (2010) was inferred from a Go/No-Go task that reflects behavioral rather than cognitive inhibition, which could explain the difference with these results; future research could investigate response inhibition more precisely to examine its role in physical endurance. Second, it is possible that the use of a cognitive task in our study reduced the importance of inhibition. Even if the duration of the endurance task was designed in such a way so as not to cause cognitive fatigue that could influence muscular endurance time, people in a physical endurance context do not usually have another task to focus on and their attention is therefore more focused on the exercise, even more so if the task induces pain (Van Damme et al., 2008). While asking participants to perform the flanker task during exercise was an excellent way to obtain an online index of their inhibition capacity, it could have also distracted participants from pain and fatigue, thus reducing the need to inhibit these signals. Future work could use an inhibition index that interferes less with attentional focus, such as monitoring brain potentials that are directly associated with inhibition (Klimesch et al., 2007)

## 5. Conclusion

To conclude, we found no effect of stimulation of the dlPFC on endurance or cognitive inhibition, despite the use of a well-controlled tDCS protocol. This lack of an effect prevented us from testing the role of cognitive inhibition in physical endurance. However, the absence of a correlation between cognitive inhibition and physical endurance suggests that this function has a limited contribution in the maintenance of physical effort. That said, it is possible that other forms of inhibition or other PFC functions play an important role in the maintenance of physical effort. We therefore encourage future research to continue this line of investigation using a neurocognitive focus, which could have an important impact in our understanding of physical fatigue.

### Ethical approval

All procedures performed in studies involving human participants were in accordance with the ethical standards of the institutional and/or national research committee and with the 1964 Helsinki declaration and its later amendments or comparable ethical standards. All persons gave their consent prior to their inclusion in the study.

